# Genetic variants disrupt human RGS_14_ binding to NHERF_1_ and regulation of NPT_2_A-mediated phosphate transport

**DOI:** 10.1101/540781

**Authors:** Peter A. Friedman, Tatyana Mamonova, Clara E. Magyar, Katherine E. Squires, W. Bruce Sneddon, David R. Emlet, John R. Hepler

## Abstract

RGS14 is a multifunctional scaffolding protein that integrates G protein, MAPK, and Ca^++^/CaM signaling pathways. RGS14 contains an RGS domain, tandem Ras/Rap-binding domains, and a G protein regulatory motif. Human/primate RGS14 differ from rodent proteins by possessing a carboxy-terminal extension encoding a Type I PDZ ligand. RGS14 has been implicated in disordered phosphate metabolism. The human *RGS14* gene is adjacent to *SLC34A1* that encodes the NPT2A sodium-phosphate cotransporter. Hormone-regulated NPT2A requires the PDZ protein NHERF1 which contains two PDZ domains (PDZ1 and PDZ2). NHERF1 binds the PDZ ligand carboxy tail of NPT2A to regulate phosphate uptake, and this NPT2A:NHERF1 complex is inhibited by parathyroid hormone (PTH). Studies here define roles for RGS14 in NHERF1-dependent, PTH-sensitive phosphate transport. We found that RGS14 binds to NHERF1 via the PDZ2 domain. PTH inhibits NPT2A-mediated phosphate transport and RGS14 blocked this action. Several rare human mutations have been reported in the RGS14 PDZ ligand located at residues 563 (D563N, D563G) and 565 (A565S, A565V). D563N disrupted RGS14 binding to NHERF1 and did not interfere with PTH action, whereas D563G, A565S, and A565V bound NHERF1 and were functionally equivalent to wild-type RGS14. Computational analysis and molecular dynamics modeling of NHERF1 PDZ2 binding to the RGS14 C-terminal PDZ ligands refined the structural determinants of this interaction. Additional studies demonstrated that RGS14 is expressed in human kidney proximal and distal tubule cells. Together, our findings are consistent with the view that RGS14 contributes to PTH-sensitive phosphate transport in humans. RGS14 coding variants may cause disordered phosphate metabolism.

---

Regulators of G protein signaling (RGS) are GTPase activating proteins (GAPs) that function primarily as central components of GPCR and G protein signaling pathways (1,2). One such RGS protein, RGS14, is an unusual multifunctional scaffolding protein that integrates G protein, MAPK, and Ca^++^/CaM signaling pathways (3,4). In rodents, RGS14 is expressed in brain, lung, and spleen (5). Most of what we know about RGS14 comes from studies in rodents, where it acts in brain as a natural suppressor of synaptic plasticity and hippocampal-based learning (6,7) and in heart to suppress cardiac remodeling (8). However, much less is known about RGS14 in humans. Human and rodent RGS14 share a common domain structure that includes an N-terminal RGS domain that binds Gαi/o-GTP acting as a GAP to limit G protein signaling (5,9), two tandem Ras/Rap-binding domains (RBD) that bind active H-Ras and Rap2 (10–12), and a G protein regulator (GPR, also known as a GoLoco) motif that binds inactive Gαi1/3 to anchor RGS14 at membranes (13). However, human/primate RGS14 differ from the rodent protein in that they also contain a C-terminal Class I PDZ-recognition sequence. Numerous genome-wide association studies have implicated RGS14 in kidney diseases (14–19), including disordered phosphate metabolism. The *RGS14* gene on human chromosome 5 is adjacent to *SLC34A1* that encodes the NPT2A sodium-phosphate cotransporter. Coding mutations have been identified (20) in the human RGS14 PDZ-binding motif, as we reported (21), though the impact of these variants on RGS14 function is unknown.

PDZ proteins, named for the common structural domain shared by the post-synaptic density protein (PSD95), Drosophila disc large tumor suppressor (DlgA), and zonula occludens 1 protein (ZO1), constitute a family of 200-300 members (22,23). These adapter molecules assemble a variety of membrane-associated proteins, including transporters, receptors, and ion channels, and signaling molecules in short-lived functional units. PDZ modules consist of 80-90 amino acids forming a 3-dimensional globular domain that is composed of six β-sheets (ßA-ßF) and 2 α-helices (αA, αB) within the larger protein (22). Scaffolding proteins harboring PDZ domains may contain single or multiple PDZ modules, and may also include other protein-protein interaction motifs (23). In the target partner protein, a C-terminal sequence, the so-called PDZ-binding motif or ligand, binds in an extended groove of the PDZ domain between the second ß-sheet (ßB) and the second α-helix (αB) in an anti-parallel fashion with the terminal hydrophobic amino acid of the ligand occupying the elongated hydrophobic cavity at the top of the binding groove. Class I PDZ ligands take the form D/E-S/T-X-Φ, where X is promiscuous and Φ is a hydrophobic residue. This sequence in human RGS14 is –DSAL.

NPT2A is expressed at luminal membranes of proximal kidney tubules, where it mediates the bulk of phosphate absorption from the urine (24). NPT2A activity is regulated by PTH and by FGF23 and requires the PDZ protein NHERF1 for this action (25). In the absence of NHERF1 or in the presence of mutations, hormone-sensitive phosphate transport is blunted with resulting hypophosphatemia (26,27). NPT2A itself contains a carboxy-terminal PDZ motif that mediates its binding to NHERF1. NHERF1 possesses two tandem PDZ domains (PDZ1, PDZ2) with identical core-binding sequences and a carboxy-terminal ezrin-binding domain, through which interacting proteins are tethered at the plasma membrane and to cortical actin (28). Recent analysis disclosed both sites upstream of the PDZ ligand and 3-dimensional organization of the PDZ module that impart greater specificity to the recognition and binding determinants between the PDZ protein and the targeted partner (29).

Based on these considerations we speculated that RGS14, acting through its PDZ-ligand, binds NHERF1 and alters PTH-sensitive phosphate transport. We theorized that the identified RGS14 variants might not share this behavior and, at the same time, afford further insight in to the structural determinants for binding of the NHERF1 PDZ domains. The goal of the work described here was to determine, for the first time, the effect of wild-type RGS14 and the identified PDZ ligand mutations on basal and PTH-regulated phosphate transport, and the role of RGS14 interactions with NHERF1 in this process. We also describe RGS14 expression in human kidney and the suppression of PTH action on phosphate transport in human kidney cells expressing wild-type RGS14.

## Results

### RGS14 binding to NHERF1

Human RGS14 contains a canonical Class I carboxy-terminal PDZ ligand consisting of the sequence –DSAL. Human NHERF1 contains two PDZ domains, PDZ1 and PDZ2 (Fig. 1*A*). To determine if RGS14 binds the PDZ protein NHERF1 we generated a FLAG-tagged RGS14 construct and performed coimmunoprecipitation experiments using HEK293 cells cotransfected with FLAG-RGS14 and HA-NHERF1. The results shown in Fig. 1*B* support the view that RGS14 binds NHERF1. We next inquired if RGS14 exhibited domain selectivity for binding with PDZ1 or PDZ2. For these experiments, we used NHERF1 constructs wherein the GYGF core-binding motif was mutated to GAGA in PDZ1 (S1), PDZ2 (S2), or both (S1/S2) PDZ domains. As shown in Fig. 1*C*, RGS14 selectively binds PDZ2.

**Figure 1.**
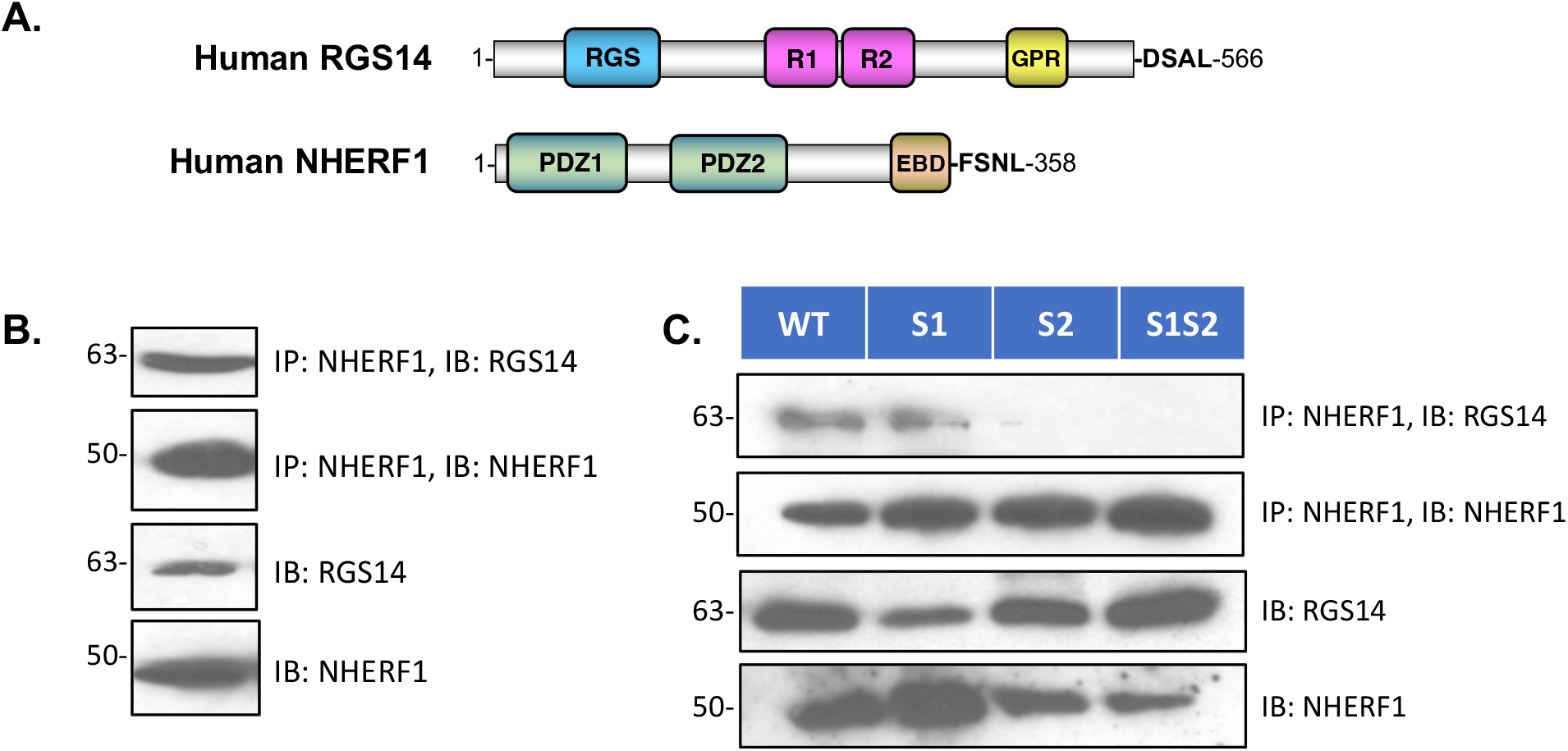
RGS14 binding to NHERF1. *A*, Diagram of the human RGS14 and NHERF1 sequence and domain structures. *B*, Coimmunoprecipitation of FLAG-RGS14 with HA-NHERF1. HEK293 cells were transfected with FLAG-RGS14 and HA-NHERF1 and recovered in complex as outlined in Experimental Procedures. *B*, RGS14 binds preferentially to the NHERF1 PDZ2 domain. HEK293 cells were cotransfected with wild-type (WT) NHERF1, or with constructs in which the core-binding motif was scrambled in PDZ1 (S1), PDZ2 (S2) or both PDZ1 and PDZ2 (S1S2) along with RGS14. Each of these experiments (*A* and *B*) was performed twice with comparable results.

Several rare coding mutations have been noted in the PDZ-binding motif of human RGS14 (21). These mutations are situated at residues D563 and A565, corresponding to positions −1 and −3 of the PDZ ligand. The specific mutations are D563N, D563G, A565S and A565V. The normal sequence and the identified mutations are summarized in Fig. 2*A*. To determine the effect of these mutations on binding to NHERF1 we prepared the various mutations in the context of full-length FLAG-tagged RGS14. Wild-type (WT) or the indicated RGS14 mutant construct was cotransfected with HA-NHERF1 in HEK293 cells. Coimmunoprecipitation experiments were performed as described in Experimental Procedures. A representative immunoblot displaying the findings for Asp^563^ is illustrated in Fig. 2*B*. The results show that replacement of Asp by Asn did not support binding to NHERF1, whereas substitution by Gly maintained the interaction with NHERF1. Compiled results are shown in Fig. 2*D*.

**Figure 2.**
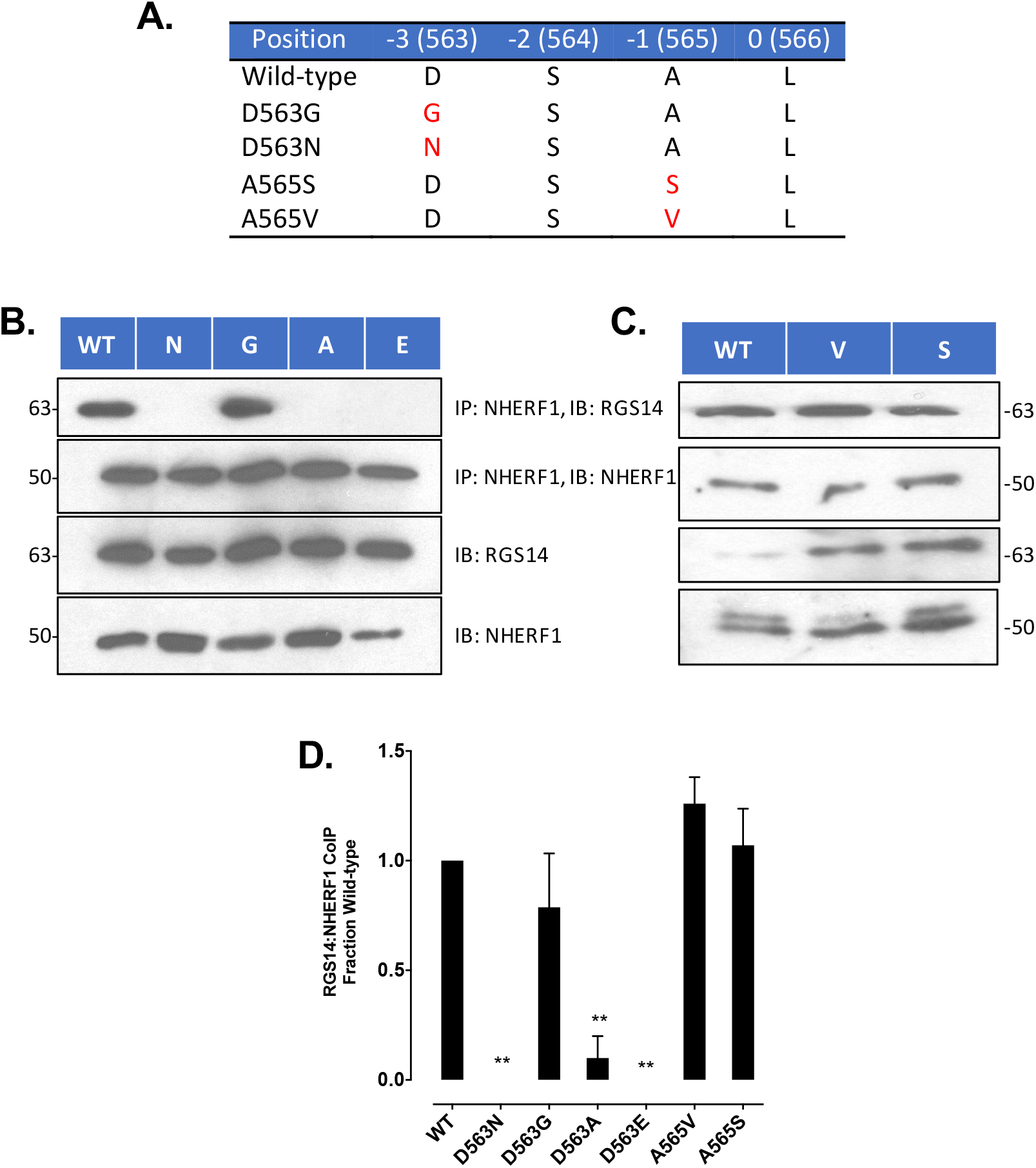
Genetic variants within the PDZ ligand of human RGS14 disrupt NHERF1 binding. *A)* Naturally occurring genetic variants and engineered mutations placed within the PDZ ligand of human RGS14. Single letter amino acid code for residues forming the C-terminal PDZ recognition motif of human RGS14. The wild-type sequence is –DSAL. Naturally occurring variants G (Gly) and N (Asn) are found at position 563, corresponding to the −3 site of the ligand, and Ser (S) and V (Val) at 565, conforming to the −1 position of the ligand. *B*, Analysis of amino acid specificity at Asp^563^ RGS14. Coimmunoprecipitation analysis of NHERF1 binding to wild-type (Asp^563^) RGS14, or with naturally occurring variants Asn (N) or Gly (G); or engineered mutants Ala (A) or Glu (E). Only Glycine supported binding with NHERF1. Similar results were observed in *n*=4 independent experiments. *C*, characterization of binding determinants at RGS position 365. Ala^565^ is a residue located at the permissive −1 position of the wild-type RGS14 PDZ ligand. Replacement with variant forms Val (V) or Ser (S) robustly coimmunoprecipitated with NHERF1. The blot is representative of 5 analyses. *D*, quantitative summary of coimmunoprecipitation experiments in *B* and *C*.

Based on these considerations, we predicted that because of its greater length Ala could not replace Asp but that by virtue of its retained charge, Glu would substitute for Asp. As shown in Fig. 1*B*, Ala^563^ failed to support interaction with NHERF1, as expected. Surprisingly, the Glu^563^ construct was unable to bind NHERF1.

We then turned our attention to Ala^565^. This corresponds to the promiscuous −1 position of the PDZ ligand (23) and hence should accommodate the reported Ser or Val variants. The results shown in Fig. 2*C* bear out this supposition inasmuch as both Ser^565^ and Val^565^ constructs avidly bound NHERF1. A quantitative summary of the coimmunoprecipitate results in given in Fig. 2*D*.

### RGS14 Regulates PTH-sensitive Phosphate Transport

Phosphate transport by the kidney is primarily mediated by the NPT2A Na-phosphate cotransporter (24). NPT2A possesses a carboxy-terminal PDZ ligand (-TRL), through which it binds NHERF1. In the absence of NHERF1 (26) or in the presence of coding region mutations (27,30), parathyroid hormone (PTH) regulation of NHERF1 is essentially eradicated resulting in profound renal phosphate wasting. RGS14 polymorphisms have been linked to disordered phosphate homeostasis in humans (14). We thus sought to determine the effect of RGS14 on basal and PTH-sensitive phosphate transport and the consequence of the various RGS14 mutations on this process. For these studies we employed opossum kidney (OK) cells, an accepted model for PTH-regulated phosphate transport (31) or human RPTEC cells, which also exhibit PTH-sensitive phosphate transport (32).

As shown in Fig. 3*A*, PTH inhibited phosphate transport by 60% in cells transfected with empty vector (EV). Hormone-sensitive phosphate transport was abolished in cells transfected with wild-type (WT) RGS14. Likewise, the RGS14 Gly^563^ mutant, which like WT interacted with NHERF1, similarly eliminated PTH-dependent phosphate transport. In contrast, the naturally occurring Asn^163^ mutation disrupted this regulatory activity, as did the artificial mutations Ala^163^ and Glu^163^. Notably, mutations that interfered with RGS14 binding to NHERF1 in turn hampered PTH-regulated phosphate transport. In contrast, the tolerated Ala^565^ variants at the permissive −1 position of the PDZ-recognition motif behaved like wild-type RGS14 and abolished PTH inhibition of phosphate uptake (Fig. 3*B*).

**Figure 3.**
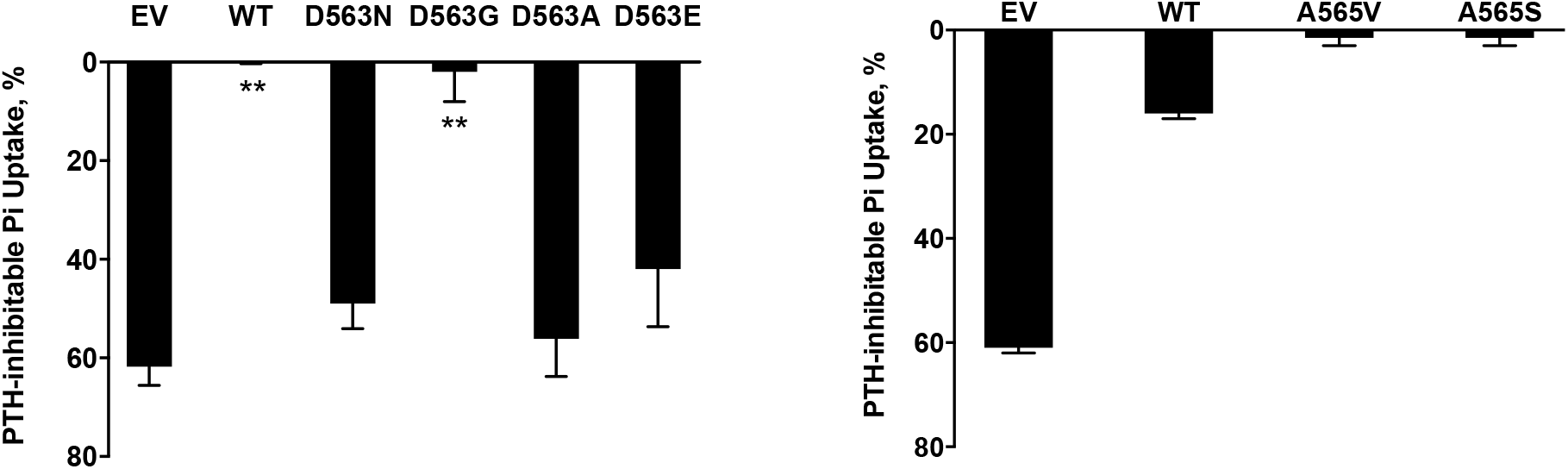
RGS14 inhibition of PTH-sensitive phosphate transport in kidney cells is disrupted by genetic variants. Phosphate transport was measured in OKH cells transfected with empty vector control (EV), wild-type NHERF1 (WT), or one of the indicated mutations at amino acids 563 (*A*) or 565 (*B*) within the human RGS14 sequence. Measurements were performed in triplicate. *n*=3-5 independent determinations. ^**^ p< 0.01.

### Kidney Expression of RGS14

The *RGS14* gene is located on chromosome 5, adjacent to *SLC34A1*, which encodes the NPT2A protein, a renal phosphate transporter. Therefore, we examined the expression of RGS14 in human kidney samples and in proximal and distal tubule cells isolated from these samples (33). As shown in Fig. 4*A*, RGS14 is expressed in protein lysates of proximal (lane 1) and distal tubule cells (lane 2). A positive control of HEK cells transfected with hRGS14 is shown in lane 3. RGS and NHERF1 colocalize in human proximal tubule sections (Fig. 4*B*) and coimmunoprecipitate in cell lysates prepared from human proximal tubule cells (Fig. 4*C*).

**Figure 4.**
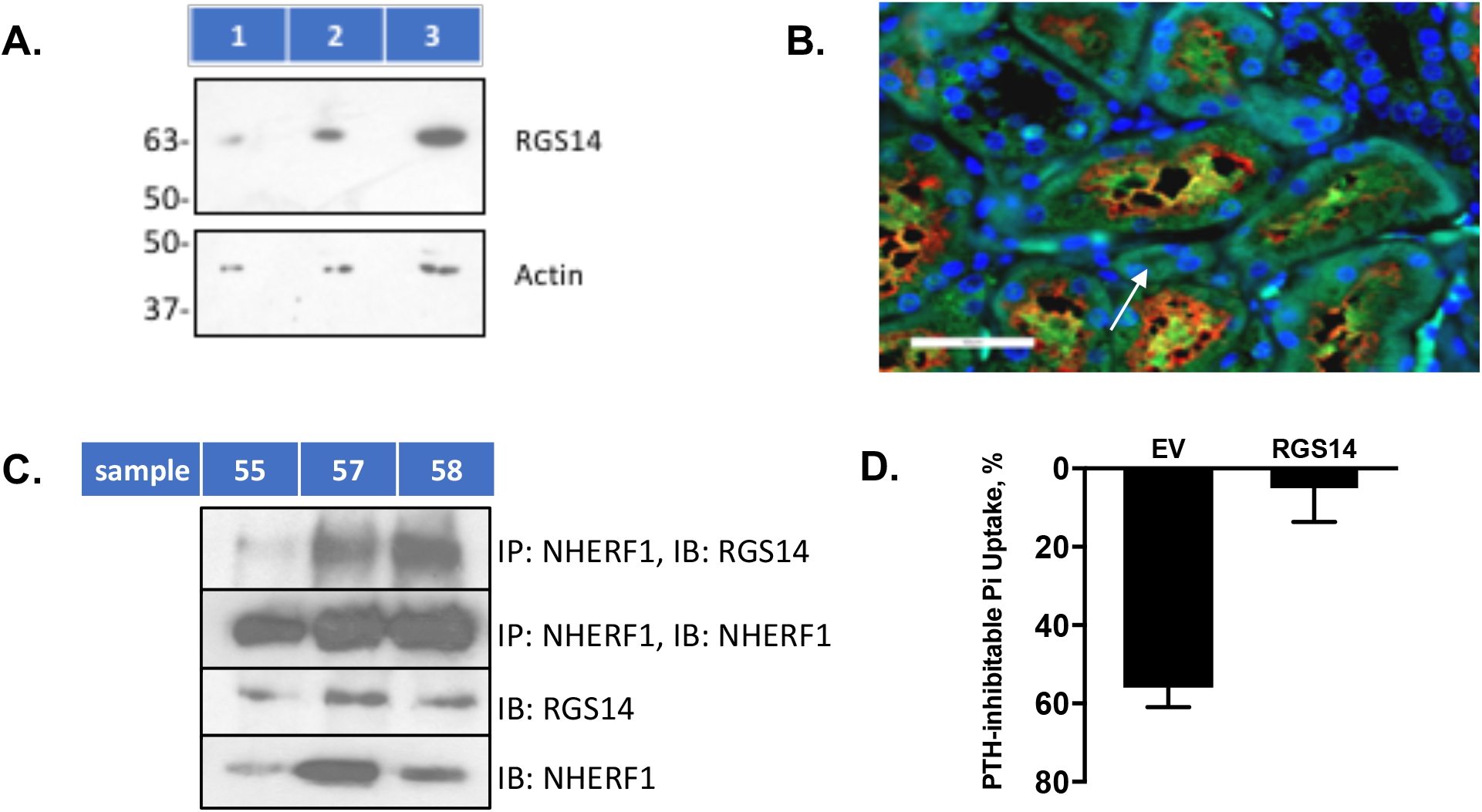
Endogenous RGS14 is expressed in human kidney and blocks PTH-sensitive phosphate uptake in human kidney cells. *A*, RGS14 in human kidney proximal (lane 1) and distal (lane 2) tubule cells and HEK cells transfected with RGS14 (lane 3). Lysates were prepared and immunoblotted as described in Experimental Procedures. Similar results were obtained in 2 experiments. *B*, RGS14, NHERF1 expression in human kidney. Representative image from whole-slide scan (40×) of formalin fixed, paraffin embedded 5-μm section of human kidney stained with RGS14 (green) and NHERF1 (red). Proximal tubular colocalization is observed (white arrow). Scale bar = 50 µm. *C,* RGS14 immunoprecipitates with NHERF1 in proximal tubule cells. NHERF1 was immunoprecipitated from donor samples 55, 57, and 58. RGS14 and NHERF1 were analyzed as detailed in Experimental Procedures. The result is illustrative of 2 experiments. *D*, RGS14 expression inhibits PTH-regulated phosphate transport in RPTEC cells. Cells were transfected with empty vector (EV) or FLAG-RGS14. 48 h later, cells were treated for 2 h with 10 nM PTH(1–34). Phosphate uptake was measured as detailed in Experimental Procedures. Data represent the mean ± S.E. (error bars) of n = 3 independent experiments performed in triplicate. Data were normalized for each experiment, where phosphate uptake under control, untreated conditions, was defined as 0% inhibition.

To extend these findings, we sought to determine and characterize the action of RGS14 on PTH-sensitive phosphate transport in RPTEC, a non-transformed line of human proximal tubule cells (34). PTH inhibits phosphate transport in RPTEC (32). Our initial experiments failed to disclose an effect on PTH action. Further examination revealed that RPTEC do not detectably express RGS14 at the message or protein level (data not shown). Accordingly, we used RPTEC as a null background to examine RGS14 actions. RPTEC were transfected with RGS14 or empty vector and PTH-sensitive phosphate transport was measured 48 h later. Inclusion of RGS14 virtually obliterated PTH action (Fig. 4*D*).

### Computational Predictions of NHERF1 PDZ2 binding to RGS14 C-terminal PDZ ligand

Key residues involved in the binding of the C-terminal peptide of RGS14 [-^558^LNSTTDSAL^566^] and PDZ2 were evaluated from explicit-solvent MD simulation. A representative structure of the PDZ2-RGS14ct-9 complex at the end of 150-ns MD simulation is shown in Fig. 5*A*. The structural stability for the complex was assessed by the root-mean-square deviation (RMSD) over the backbone atoms of the complex, PDZ2 and the RGS14ct-9 peptide. The RMSD values stabilized after approximately 50-ns (not shown). The average RMSD values along the last 20-ns of MD trajectory are 2.1 ± 0.1 Å, 1.7 ± 0.1 Å and 1.8 ± 0.1 Å for the complex, PDZ2, and RGS14ct-9, respectively. These values are relatively small and indicate that the resulting complex is structurally stable during the MD simulations. Visual analysis of the evaluated structures confirmed that the docking position of the C-terminal RGS14ct-9 peptide is similar to that of other PDZ-ligand systems (29,35-37). The RGS14ct-9 C-terminus occupies the binding groove between the α2 helix and β2 sheet, whereas Leu-0 engages deep in the hydrophobic binding pocket making canonical interactions with the residues from the carboxylate-binding loop of PDZ2 (-^163^GYGF^166^-), α2 helix (Val^216^ and Arg^220^) and β2 sheet (Leu^170^). Another canonical interaction is formed between the OH group of Ser-2 and His^212^. The distance between the carboxylate group of Asp-3 and the α-amino group of Arg^180^ or Asn^169^ stabilizes between 2-6 Å with an average of 4.5 Å and 3.7 Å for the Asp-3 Oδ1 and Arg^180^ Nη22 and Asp-3 Oδ2 and Arg^180^ Nη12, respectively, and 4.6 Å for the Asp-3 Oδ1 and Asn^169^ Nη21 (Fig. 5*A*). These ionic pairs may play a critical role in PDZ2-RGS14 association. In addition, the upstream Thr-4 is close enough to Gly^170^ to form a backbone hydrogen bond.

**Figure 5.**
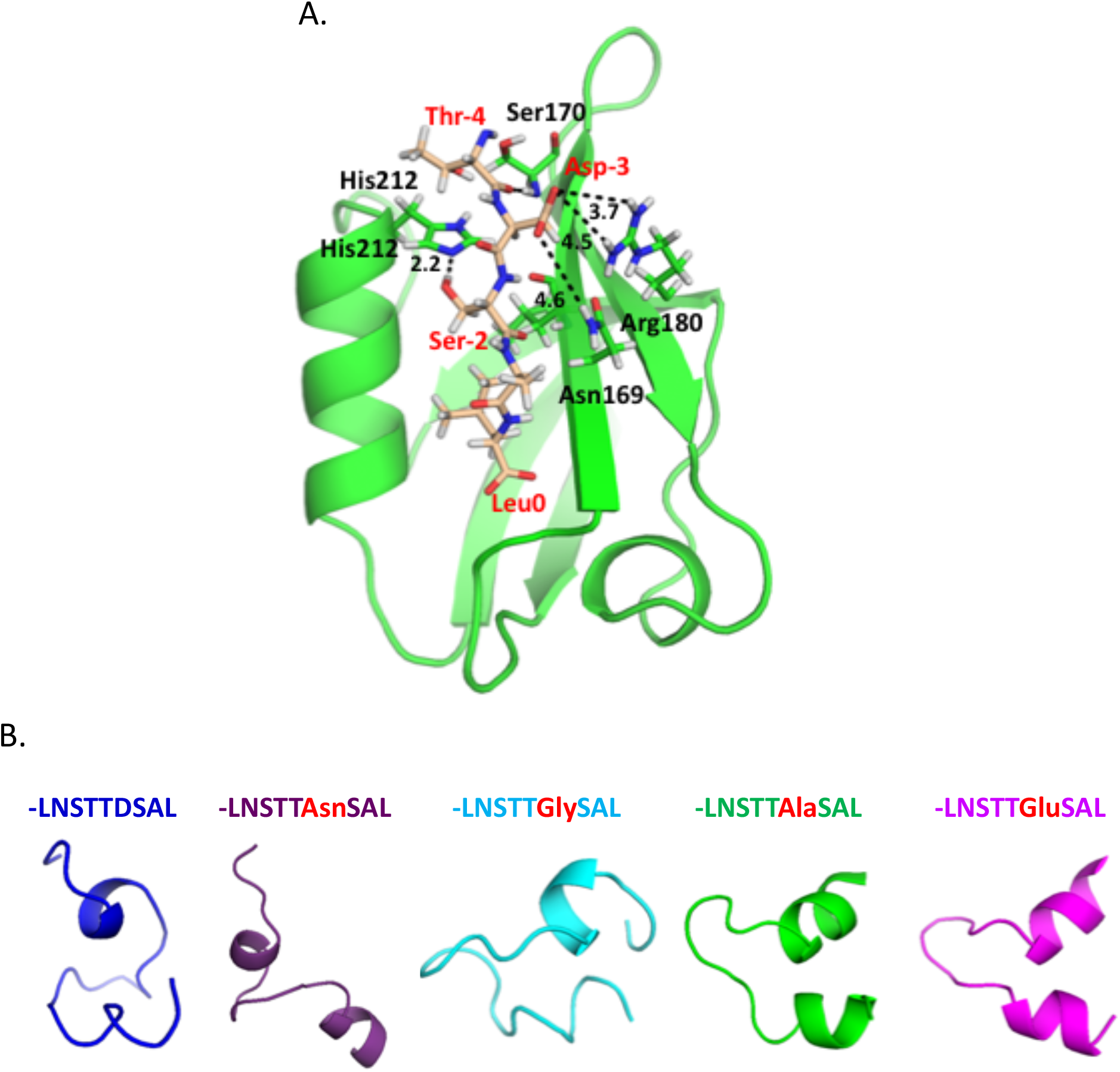
Computational prediction of NHERF1 PDZ2 binding to RGS14 C-terminal PDZ ligand and modeling of RGS14 coding variants. *A,* model structure of PDZ2 bound the C-terminal peptide of RGS14ct-9. PDZ2 is highlighted in green cartoon, whereas the RGS14ct-9 peptide is represented in wheat sticks. The last four N-terminal residues of the peptide not involved in the interaction with PDZ2 are omitted for simplicity. The key residues (except the carboxy-terminal motif) stabilizing the complex are highlighted and labeled on the structure. The dotted line represents a hydrogen bond or salt bridge between a hydrogen atom and acceptor with a distance labeled in Å. Hydrogen atoms are white, oxygens are red, and nitrogens are blue. *B,* representative structures of the C-terminal peptides of WT and mutated RGS14. The structures of the unbound 23 residues peptide of RGS14 with Asn, Gly, Ala or Glu at the position −3 (purple, cyan, green and magenta, respectively) has a tendency to form a helical segment at the C-terminus while a peptide with the Gly substitution (cyan) has a similar fold as WT RGS14ct-9 (blue). The structures were generated using Pep-FOLD3 on-line computational framework.

### Modeling of RGS14 Coding Variants

The binding to PDZ2 of four C-terminal peptides corresponding to the described mutations in RGS14 (Asp^563^Asn, Asp^563^Gly, Asp^563^Ala and Asp^563^Glu) was tested using MD simulation (Fig. 5*B*). The wild-type [PDZ2-RGS14ct-9] complex was used as a template to construct a model for each mutant, where Asp-3 in RGS14ct-9 was replaced iteratively by Asn, Gly, Ala and Glu. Because canonical PDZ-ligand interactions occur when the target peptide occupies the PDZ binding site as a short length of β-sheet (3-4 C-terminal residues), wild-type RGS14ct-9 peptide or mutated variants were modeled as linear peptides bound to PDZ2. 70-ns MD trajectories were generated for each mutated complex except 150-ns for the Gly mutant (for details see Experimental Procedures). A standard PDZ domain topology was observed for all structures, with the C-terminal binding motif positioned in an extended position between the α2 helix and β2 sheet of PDZ2. Despite the residue differences at positon −3 of the C-terminal motifs, we did not observe significant deviations from the naturally occurring Asp. Contrary from what was predicted from the experimental data (Fig. 1*C*) RGS14ct-9Asn, RGS14ct-9Ala, and RGS14ct-9Glu participate in the canonical contacts specific for Class I motifs (-D/E-S/T-X-Φ). We hypothesize that the substitution by Asn, Ala, or Glu at Asp-3 induces conformational changes or promotes the formation of local secondary structure at the C-terminal motif of RGS14, thus, preventing association with the PDZ2 domain. To test this theory we applied PEP-FOLD3 (38) to predict the structure of the C-terminal fragment (23 amino acid residues) of wild-type RGS14 and the corresponding mutants. The five best models for wild-type and mutated peptides were acquired and viewed using PyMol (Molecular Graphics System, Version 2.0 Schrödinger, LLC). Representative structures of the peptides are shown in Fig. 5*B*. As can be seen, replacement of Asp-3 by Asn, Ala, or Glu promotes the formation of a helical motif at the C-terminus, whereas the peptide with Gly is similar to wild-type (Asp) and does not demonstrate this tendency. These findings provide provisional support for this theory and a starting point for further investigation and refinement.

## Discussion

The goal of these studies was to define a possible role for RGS14 in hormone-sensitive phosphate transport. The results described here introduce novel protein-protein interactions that participate in this activity. We report here for the first time that human RGS14 directly binds NHERF1 to form a stable complex, and that this RGS14:NHERF1 complex blocks PTH-sensitive phosphate uptake by NPT2A in renal proximal tubule cells (Fig. 6). Our findings also specify the structural determinants for these interactions. The findings show for the first time that RGS14, a GAP for Gαi/o proteins, is expressed in human kidney, and more specifically in proximal and distal tubules. Interestingly, the –DSAL PDZ ligand in human RGS14 is not well conserved and is absent in the rodent protein. Thus, the actions, and the neural, cardiovascular phenotype of *RGS14* knockout mice (6,8), cannot be attributed to the absence of interactions with a PDZ partner protein. The RGS14 PDZ ligand is present throughout the primate order and in sheep. RGS14 has been implicated by numerous GWAS studies in various forms of chronic human kidney diseases (14–19). Relevant to that, we identify here one human genetic variant (D563N) that disrupts RGS14 binding to NHERF1 and its capacity to inhibit PTH-sensitive phosphate uptake.

**Figure 6.**
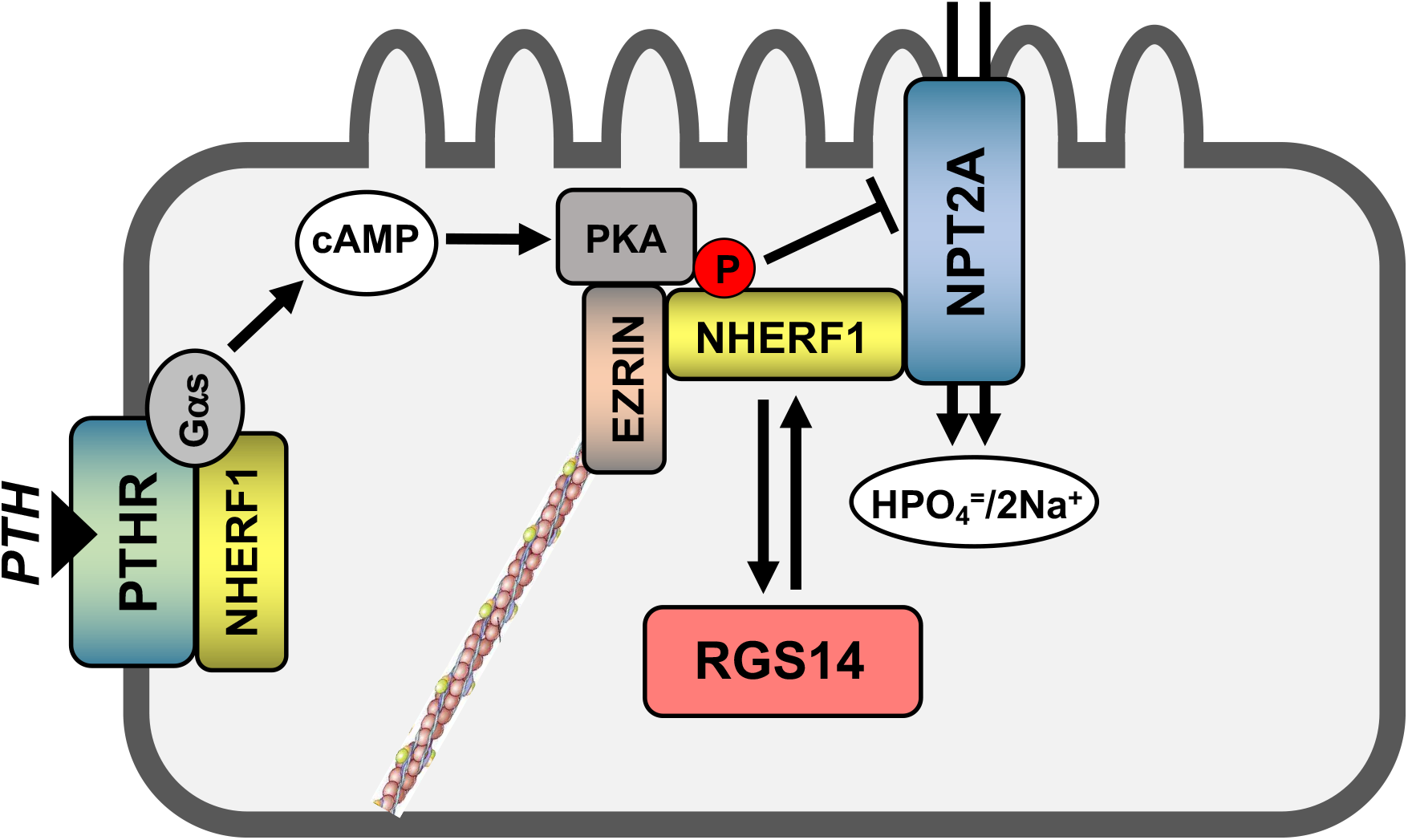
Proposed model for RGS14 regulation of PTH-sensitive phosphate uptake by NPT2A in renal proximal tubule cells. The [NPT2A:NHERF1:Ezrin] protein complex promotes uptake of phosphate across apical membrane renal proximal tubule cells. PTH activation of the PTH receptor (PTHR)-Gs complex stimulates adenylyl cyclase to generate cAMP, which activates PKA bound to Ezrin. PTH-induced activation of PKA, along with PKC, and GRK6 (not shown) phosphorylate NHERF1 to uncouple it from NPT2A and inhibit phosphate uptake. Findings described here show that the C-tail PDZ-ligand of RGS14 binds the PDZ2 domain of NHERF1 to block PTH-induced inhibition of NPT2A phosphate uptake. Furthermore, we show that human genetic variants within the PDZ ligand of RGS14 disrupt RGS14 actions on NHERF1 and PTH-sensitive phosphate uptake.

The initial results affirmed that RGS14 binds NHERF1 and, more specifically to PDZ2. The general motif of a 4-part type 1 PDZ ligand takes the form D/E-S/T-X-Φ. The particular RSG14 ligand sequence is -AspSerAlaLeu. Based on these considerations we expected that Asn or Gly mutations at Asp-3 (Asp^563^) would disrupt binding to NHERF1. Although Asn substitution abolished binding as predicted, Gly replacement was innocuous. Asn contains an amide group in place of one of the Asp carboxyl groups, making Asp electroneutral. Gly, in contrast, is shorter and contains a proton as its side-chain, giving it greater conformational flexibility. Thus, in this instance, Gly^563^ is permissive for binding to NHERF1. A related conundrum emerged when we examined Ala and Glu substitutions at Asp^563^. Here, we predicted that Ala could not replace the shorter Gly, and that negatively charged Glu would be a surrogate for the naturally occurring, negatively charged Asp. This too turned out to be partially incorrect inasmuch as Glu could not replace Asp, again, a difference of a single carbon group. Based on these structural constraints, we performed Molecular Dynamics (MD) simulations to model the residue-specific interactions of the RGS14 PDZ ligand with NHERF1 PDZ2.

The C-terminus RGS14 PDZ-recognition motif (-DSAL) is a permissive sequence that in principle could interact with both NHERF1 PDZ1 and PDZ2 domains, which have extensive similarity and share identical GYGF-core binding motifs (35). Despite the fact that PDZ1 interacts with a larger and more diverse set of ligands compared to PDZ2 (39–42), the latter specifically recognizes the C-terminal motif of RGS14. In the absence of experimental structural information, we developed a computational model, where PDZ2 binds the 9 residues C-terminal motif of RGS14. The model we advance here suggests that in addition to the canonical RGS14 PDZ ligand interactions between Leu-0 and the residues from the -^163^GYGF^166^-loop of PDZ2, and Ser-2 and His^212^, Arg^180^ from the β3 strand and Asn^169^ from the β2 strand further stabilize the PDZ2-RGS14 complex. The predicted electrostatic interactions are similar to the Arg^40^/Arg^180^-Glu-3 ionic pair observed for association of PDZ1/PDZ2 with PTHR as reported (43). Although previous work highlights the importance of residues in upstream sequences promoting the coordinated interaction of peptides to the extended binding groove (37,44), upstream Thr-5, Ser-6, Asn-7 and Leu-8 of RGS14 are not involved in forming stable interactions with PDZ2 on the MD simulation time scale.

MD simulations performed for PDZ2 bound peptides with the Asp-3/Asn-3, Asp-3/Gly-3, Asp-3/Ala-3 and Asp-3/Glu-3 substitution revealed that the alternate peptides occupied the PDZ2 domain binding groove in an orientation and conformation similar to the wild-type RGS14. These predictions were contrary to the findings predicted from the biochemical experiments, where the single Asp-3/Asn-3, Asp-3/Ala-3, Asp-3/Glu-3 but not Asp-3/Gly-3 mutation destabilized the interaction with NHERF1. The function of the C-terminus of RGS14 is related to its flexibility and ability to occupy the binding pocket of the PDZ2 domain. We theorized that Glu or Asn variants with the longer side chains compared to Asp-3, or the Ala variant with a methyl group as a side chain, but not Gly, could promote distinct conformational changes of the C-terminal motif of RGS14. Using minimalist computational modeling, we showed that the charged (Glu), polar (Asn) or hydrophobic (Ala) patterns in the RGS14 C-terminal peptide can promote formation of a helix. Modeling of the peptide with the Asp-3/Gly-3 substitution revealed that the peptide is structurally stable and remains in the conformation similar to the wild-type RGS14 C-terminal motif. Based on this observation we reasoned that RGS14ct-9Gly could bind PDZ2 in the canonical PDZ-ligand mode. Structures evaluated from MD simulations showed high similarity between the orientation of Leu-0 and Ser-2 of PDZ2-RGS14ct-9 and PDZ2-RGS14-9Gly. A notable difference between the two peptide ligands is observed at the N-terminal end.

We speculate that loss of the electrostatic interaction between Asp-3/Gly-3 and Arg^180^ promotes conformational changes upstream of ligand position −3. Thr-4 changes its position and orientation to maintain a hydrogen bond equivalent to the Asp-3-Arg^180^ interaction in wild-type PDZ2-RGS14ct-9. Formation of new specific contacts between Thr-4 with Asn^169^ and Arg^180^ might stabilize the PDZ2-RGS14ct-9Gly complex. We propose that distinct conformational changes in the C-terminal motif of RGS14 induced by the mutations, except Gly, which is tolerated, impede association between PDZ2 and RGS14, and the functional role of Asp-3 is to support a dynamic network between average conformations of the RGS14 C-terminus and its ability to occupy the PDZ2 binding pocket and interact with PDZ2

Although RGS14 has been studied in a variety of tissues, its relevance to PTH-regulated phosphate transport by NPT2A would require kidney expression, which had not been described. We found that RGS14 is well expressed in human kidney and in isolated proximal and distal tubule cells. Notably, NPT2A is found exclusively in proximal tubules (45). Hence, the presence of RGS14 in distal tubules cells relates to some independent regulatory activity.

In summary, we report here for the first time that human RGS14 plays an important role in PTH-regulated phosphate transport by NPT2A, and that this regulatory role of RGS14 involves direct and specific binding of its cognate PDZ ligand to the PDZ2 domain of NHERF1 (Fig. 6). We determined the effects of identified human genetic variants within the PDZ motif of RGS14 on NHERF1 binding, and found that one (Asp^563^Asn) disrupted the RGS14:NHERF1 complex and blocked RGS14 actions. RGS14 is implicated by numerous GWAS studies to be involved in various forms of chronic kidney diseases (14–19). Relevant to that, we identified here at least one human genetic variant that blocks RGS14 actions on PTH-sensitive phosphate uptake. Ongoing and future studies will further define novel signaling roles for RGS14 in proximal and distal renal tubules in both physiology and disease.

### Experimental Procedures

#### Chemical Reagents, Plasmids and Antibodies

Monoclonal anti-HA agarose (A2095) and rabbit polyclonal anti-FLAG (F7425) were purchased from Sigma. Rabbit polyclonal anti-HA (sc-805) and protein G+ agarose (sc-2002) were purchased from Santa Cruz Biotechnology. Rabbit polyclonal anti-NHERF1 was purchased from Abcam (ab3452). Rabbit polyclonal anti-RGS14 was purchased from Proteintech (16258-1-AP). HA-NHERF1 was generated as described (46). [Nle^8,18^,Tyr^34^]PTH(1–34) was from Bachem, Torrance, CA (H9110).

#### Human Genetic Variants and Constructs

Human variants of RGS14 were acquired from the Genome Aggregation Database (GnomAD version 2.0, Broad Institute) and filtered for missense variants. Two human variants were identified (D563G and D563N) and selected based on the change in side chain properties from charged to neutral (D563G) or to polar (D563N). Mutations were added to FLAG-RGS14 using Qiagen QuikChange II and the following primers. D563G: GGA TCC TTG AAC TCC ACC ACC GGC TCA GCC CTC (forward) and GAG GGC TGA GCC GGT GGT GGA GTT CAA GGA TCC (reverse). D563N: GGA TCC TTG AAC TCC ACC ACC AAC TCA GCC CTC (forward) and GAG GGC TGA GTT GGT GGT GGA GTT CAA GGA TCC (reverse).

#### Cell Lines and Culture

Opossum kidney cells (OK) were obtained from J. Cole (47) and cultured in DMEM/F-12 (Corning, 10-090-CV) supplemented with 5% heat-inactivated fetal bovine serum (GenClone 25-514H, Genesee Scientific) plus 1% penicillin and streptomycin. Human Embryonic Kidney cells (HEK) were obtained from ATCC. They were cultured in DMEM (Corning, 10-013-CV) supplemented with 10% FBS plus 1% penicillin and streptomycin. Human renal proximal tubule epithelial cells immortalized with hTERT (RPTEC) were obtained from ATCC under license from Geron Corp. They were cultured in defined medium (DMEM/F-12 (Corning, 10-090-CV) supplemented with 5 pM triiodo-L-thyronine, 10 ng/ml recombinant human epidermal growth factor, 25 ng/ml prostaglandin E1, 3.5 µg/ml ascorbic acid, 1 mg/ml insulin, 0.55 mg/ml transferrin, 0.5 µg/ml sodium selenite, 25 ng/ml hydrocortisone) plus 1% penicillin and streptomycin and 0.1 mg/ml G418.

#### Tissue and Cell Preparation

Human adult kidney (HAK10, 65yo M; HAK55, 63yo/M; HAK57, 53yo/M; HAK58, 63yo/F); CD13 (proximal tubule), and CD227 (distal tubule) cells were from male and female donors, as indicated. Tissue procurement and processing have been described (33). CD13 and CD227 cells were seeded on 6 well plates. Lysates were prepared using 1% Nonidet P-40 buffer supplemented with protease inhibitor mixture I (EMD Millipore).

Protein lysates from mouse kidney were prepared by mincing tissue with a razor blade on an ice-cold metal block. The minced kidney was then homogenized using a Cole-Parmer PTFE Tissue Grinder (30 ml size) in 2 ml of ice-cold 1% Nonidet P-40 (50 mM Tris, 150 mM NaCl, 5 mM EDTA, 1% Nonidet P-40) supplemented with protease inhibitor mixture I. After incubation on ice for 30 min, the lysate was centrifuged at 13,200 rpm in an Eppendorf 5415-R refrigerated microcentrifuge. The supernatant (mouse kidney protein lysate) was saved for immunoblot analysis.

Protein lysates were prepared from HEK293 cells transfected with RGS14. Cells were seeded on 6-well plates. 24 h later, the cells were transfected with 1µg/well of human RGS14 expression plasmid using JetPrime (Polyplus). After 48 h, protein lysates were prepared using 1% Nonidet P-40 buffer supplemented with protease inhibitor mixture I.

#### Coimmunoprecipitation, Immunoblotting

HEK cells were seeded on 6-well plates. 24 h later, cells were transfected with 1µg each/well of HA-NHERF1 and FLAG-RGS14 (WT, mutants as indicated) using jetPRIME® (Polyplus, New York, NY). 48 h post-transfection, cells were lysed with 1% Nonidet P-40 (50 mM Tris, 150 mM NaCl, 5 mM EDTA, 1% Nonidet P-40) supplemented with protease inhibitor mixture I (EMD Millipore). Lysis was performed for 15 min on ice. NHERF1 was immunoprecipitated (IP) using anti-HA agarose overnight at 4 °C. IP samples were washed 4 times with 400 µl of lysis buffer and pelleted by spinning at 6000 rpm for 5 min at 4 °C in an Eppendorf 5415-R refrigerated microcentrifuge. The final pellet was resuspended in 60 µl of SDS-PAGE sample supplemented with 5% β-mercaptoethanol (Sigma). Immunoprecipitated proteins were eluted by incubating samples for 5 min at 95 °C and then pelleting the agarose by spinning for 1 min at 13200rpm in an Eppendorf 5415 microcentrifuge. Supernatant proteins were resolved on 10% SDS-polyacrylamide gels and transferred to Immobilon-P membranes (Millipore) using the semidry method (Bio-Rad). Membranes were blocked for 1 h at room temperature with 5% nonfat dried milk in Tris-buffered saline plus Tween 20 (TBST) (blocking buffer) and incubated with the primary antibodies (polyclonal anti-FLAG at 1:1000, polyclonal anti-HA at 1:1000) in blocking buffer overnight at 4 °C. The membranes were washed four times for 10 min in TBST and then incubated with goat anti-rabbit IgG conjugated to horseradish peroxidase at a 1:5000 dilution for 1 h at room temperature. Membranes were washed four times for 10 min in TBST. Protein bands were detected by Luminol-based enhanced chemiluminescence (EMD Millipore WBKLS0500).

#### Phosphate Transport

RPTEC or OK cells were seeded on 12 well plates. 24 h later, cells were transfected with 1µg/well of WT or mutant FLAG-RGS14 plasmid, as indicated, using jetPRIME (Polyplus, Strasbourg, FR). 48hr later, cells were serum-starved overnight and then treated for 2 h with 10 nM (RPTEC) or 100 nM (OK cells) PTH(1–34). Phosphate uptake was measured as described (32). The hormone-supplemented medium was aspirated, and the wells were washed three times with 1 ml of Na-replete wash buffer (140 mM NaCl, 4.8 mM KCl, 1.2 mM MgSO_4_, 0.1 mM KH_2_PO_4_, 10 mM HEPES, pH 7.4). The cells were incubated with 1 µCi [^32^P]orthophosphate (PerkinElmer Life Sciences, NEX053) in 1 ml of Na-replete buffer for 10 min. Phosphate uptake was terminated by placing the plate on ice and rinsing the cells three times with Na-free buffer (140 mM N-methyl-D-glucamine, 4.8 mM KCl, 1.2 mM MgSO_4_, 0.1 mM KH_2_PO_4_, 10 mM HEPES, pH 7.4). The cells in each well were extracted overnight at 4 °C using 500 µl 1% Triton X-100 (Sigma). A 250-µl aliquot was counted in a Beckmann Coulter LS6500 scintillation spectrometer. Data were normalized to phosphate uptake under control conditions defined as 100%.

#### Immunofluorescent staining and Image Capture

Immunofluorescent staining was performed with the PerkinElmer Opal kit (Perkin-Elmer, Waltham, MA). Briefly, 5-µm thick FFPE sections, baked and deparaffinized, were processed for antigen retrieval, blocking, primary and secondary antibody incubations and signal amplification according to manufacturer’s protocol. RGS14 (16258-1, Proteintech, Rosemont, IL, 1:200) was visualized using Opal 520 (green) and NHERF1 (EBP50, ab3452, Abcam, Cambridge, MA, 1:200) was visualized using Opal 570 (red).

Fluorescent images were acquired using the Aperio Versa Digital Pathology Scanner (Leica Biosystems, Buffalo Grove, IL). The Versa scanner is based on a Leica DM6000B microscope with motorized stage and autofocus capabilities. Slides were scanned at 40X objective magnification with the DAPI, FITC and TRITC filters. Optimal exposure times were determined before automated scanning.

#### System Modeling and Protocols for MD Simulation and Analysis

Molecular Dynamics (MD) simulations were performed using the AMBER16 package with the AMBER force field (ff99SB). The model of NHERF1 PDZ2 bound carboxy-terminal fragment of RGS14 was prepared using the PDZ2-PTHR complex as a template (43). In the Leap program (48) the 9-residue, carboxy-terminal –^585^LQEEWETVM^593^ motif of PTHR was replaced by –^558^LNSTTDSAL^566^ in RGS14. The complex was then solvated with TIP3P water molecules in a periodically replicated box, neutralized with a Na^+^ ion, and energy minimized over 500 steps including 100 steps of steepest descent minimization using the AMBER 16 pmemd module.

Equilibration and production simulations were performed as detailed previously (43). Briefly, the short runs (0.7 −0.8 ns) were conducted under the NPT ensemble [constant number of particles (N), pressure (P), and temperature (T)] to equilibrate the water molecules. During this equilibration harmonic restrains were applied to the ligand and methodically lowered from *k_s_* = 10 kcal/mol/Å^2^ to 0.1 kcal/mol/Å^2^. Then, equilibration runs (40-50 ns) were continued under the NVT ensemble [constant number of particles (N), volume (V), and temperature (T)] with harmonic restrains *k_s_* = 0.1 kcal/mol/Å^2^ applied to the N-terminal backbone atoms of the ligand and the PDZ domain. Production simulations (100 ns) were carried out with similar weak harmonic restrains (*k_s_* = 0.1 kcal/mol/Å^2^) to prevent diffusion of the complex from the simulation box. All simulations were performed at 300 K with configurations saved every 2 fs for analysis. Analysis of the trajectories was carried out with cpptraj, a complementary trajectory analysis program in the AMBER 16 suite. The RMSDs of the backbone atoms (N, CA, C) relative to their starting positions were calculated for the PDZ2-RGS14 complex, PDZ2 and RGS14ct-9 over the entire MD trajectory. The absence of the backbone conformational changes for the core of PDZ2 as well as for the carboxy-terminal motif of the bound peptide during the equilibration and production simulation indicates that the resulting complex is stable and remains close to the initial structure.

#### Statistical Analysis

Results were analyzed using Prism 7 software (GraphPad, La Jolla, CA). Data represent the mean ± SE of *n* ≥ 3 independent experiments and were compared by analysis of variance with *post hoc* testing using the Bonferonni procedure. *p* values < 0.05 were considered statistically significant.

## Acknowledgments

National Institutes of Health (NIH) awards DK105811, DK111427 (PAF); NS037112, NS102652 (JRH). The authors would like to thank Suneela Ramineni for excellent technical contributions.

## Footnotes

Human proteins are indicated by 3-letter uppercase convention; genes in italics. Only the first letter is uppercase for the corresponding mouse protein or gene.

The abbreviations used are: PTH, *parathyroid hormone*; PTHR, *PTH* receptor; NPT2A, *Sodium-phosphate cotransport protein 2A*; RGS14, *regulator of G protein signaling 14*; PDZ, *postsynaptic density protein 95 (PSD95), drosophila disc large tumor suppressor (DlgA), and zonula occludens 1 protein (ZO1)*; GAP, *GTPase activating protein*; RBD, *ras-binding domain*; GPR, *G protein regulatory domain*; PKA, *cAMP-dependent protein kinase*; RPTEC, *normal human Primary Renal Proximal Tubule Epithelial Cells*; OKH, *NHERF1-deficient Opossum Kidney cells*; MD, *molecular dynamics simulation*; HEK, *Human Embryonic Kidney cells*

## References

1. Hollinger, S., and Hepler, J. R. (2002) Cellular regulation of RGS proteins: modulators and integrators of G protein signaling. Pharmacol Rev 54, 527–559

2. Ross, E. M., and Wilkie, T. M. (2000) GTPase-activating proteins for heterotrimeric G proteins: regulators of G protein signaling (RGS) and RGS-like proteins. Annu Rev Biochem 69, 795–827

3. Evans, P. R., Dudek, S. M., and Hepler, J. R. (2015) Regulator of G Protein Signaling 14: A Molecular Brake on Synaptic Plasticity Linked to Learning and Memory. Prog Mol Biol Transl Sci 133, 169–206

4. Evans, P. R., Gerber, K. J., Dammer, E. B., Duong, D. M., Goswami, D., Lustberg, D. J., Zou, J., Yang, J. J., Dudek, S. M., Griffin, P. R., Seyfried, N. T., and Hepler, J. R. (2018) Interactome Analysis Reveals Regulator of G Protein Signaling 14 (RGS14) is a Novel Calcium/Calmodulin (Ca(2+)/CaM) and CaM Kinase II (CaMKII) Binding Partner. J. Proteome Res. 17, 1700–1711

5. Hollinger, S., Taylor, J. B., Goldman, E. H., and Hepler, J. R. (2001) RGS14 is a bifunctional regulator of Galphai/o activity that exists in multiple populations in brain. J. Neurochem. 79, 941–949

6. Lee, S. E., Simons, S. B., Heldt, S. A., Zhao, M., Schroeder, J. P., Vellano, C. P., Cowan, D. P., Ramineni, S., Yates, C. K., Feng, Y., Smith, Y., Sweatt, J. D., Weinshenker, D., Ressler, K. J., Dudek, S. M., and Hepler, J. R. (2010) RGS14 is a natural suppressor of both synaptic plasticity in CA2 neurons and hippocampal-based learning and memory. Proc Natl Acad Sci U S A 107, 16994–16998

7. Evans, P. R., Parra-Bueno, P., Smirnov, M. S., Lustberg, D. J., Dudek, S. M., Hepler, J. R., and Yasuda, R. (2018) RGS14 restricts plasticity in hippocampal CA2 by limiting postsynaptic calcium signaling. eNeuro 5

8. Li, Y., Tang, X. H., Li, X. H., Dai, H. J., Miao, R. J., Cai, J. J., Huang, Z. J., Chen, A. F., Xing, X. W., Lu, Y., and Yuan, H. (2016) Regulator of G protein signalling 14 attenuates cardiac remodelling through the MEK-ERK1/2 signalling pathway. Basic Res Cardiol 111, 47

9. Cho, H., Kozasa, T., Takekoshi, K., De Gunzburg, J., and Kehrl, J. H. (2000) RGS14, a GTPaseactivating protein for Gialpha, attenuates Gialpha-and G13alpha-mediated signaling pathways. Mol. Pharmacol. 58, 569–576

10. Traver, S., Bidot, C., Spassky, N., Baltauss, T., De Tand, M. F., Thomas, J. L., Zalc, B., Janoueix-Lerosey, I., and Gunzburg, J. D. (2000) RGS14 is a novel Rap effector that preferentially regulates the GTPase activity of galphao. Biochem J 350 Pt 1, 19–29

11. Willard, F. S., Willard, M. D., Kimple, A. J., Soundararajan, M., Oestreich, E. A., Li, X., Sowa, N. A., Kimple, R. J., Doyle, D. A., Der, C. J., Zylka, M. J., Snider, W. D., and Siderovski, D. P. (2009) Regulator of G-protein signaling 14 (RGS14) is a selective H-Ras effector. PLoS One 4, e4884

12. Shu, F. J., Ramineni, S., and Hepler, J. R. (2010) RGS14 is a multifunctional scaffold that integrates G protein and Ras/Raf MAPkinase signalling pathways. Cell Signal 22, 366–376

13. Shu, F. J., Ramineni, S., Amyot, W., and Hepler, J. R. (2007) Selective interactions between Giα1 and Giα3 and the GoLoco/GPR domain of RGS14 influence its dynamic subcellular localization. Cell Signal 19, 163–176

14. Kestenbaum, B., Glazer, N. L., Kottgen, A., Felix, J. F., Hwang, S. J., Liu, Y., Lohman, K., Kritchevsky, S. B., Hausman, D. B., Petersen, A. K., Gieger, C., Ried, J. S., Meitinger, T., Strom, T. M., Wichmann, H. E., Campbell, H., Hayward, C., Rudan, I., De Boer, I. H., Psaty, B. M., Rice, K. M., Chen, Y. D., Li, M., Arking, D. E., Boerwinkle, E., Coresh, J., Yang, Q., Levy, D., van Rooij, F. J., Dehghan, A., Rivadeneira, F., Uitterlinden, A. G., Hofman, A., van Duijn, C. M., Shlipak, M. G., Kao, W. H., Witteman, J. C., Siscovick, D. S., and Fox, C. S. (2010) Common genetic variants associate with serum phosphorus concentration. J Am Soc Nephrol 21, 1223–1232

15. Urabe, Y., Tanikawa, C., Takahashi, A., Okada, Y., Morizono, T., Tsunoda, T., Kamatani, N., Kohri, K., Chayama, K., Kubo, M., Nakamura, Y., and Matsuda, K. (2012) A genome-wide association study of nephrolithiasis in the Japanese population identifies novel susceptible Loci at 5q35.3, 7p14.3, and 13q14.1. PLoS genetics 8, e1002541

16. Yasui, T., Okada, A., Urabe, Y., Usami, M., Mizuno, K., Kubota, Y., Tozawa, K., Sasaki, S., Higashi, Y., Sato, Y., Kubo, M., Nakamura, Y., Matsuda, K., and Kohri, K. (2013) A replication study for three nephrolithiasis loci at 5q35.3, 7p14.3 and 13q14.1 in the Japanese population. J Hum Genet 58, 588–593

17. Mahajan, A., Rodan, A. R., Le, T. H., Gaulton, K. J., Haessler, J., Stilp, A. M., Kamatani, Y., Zhu, G., Sofer, T., Puri, S., Schellinger, J. N., Chu, P. L., Cechova, S., van Zuydam, N., Consortium, S., BioBank Japan, P., Arnlov, J., Flessner, M. F., Giedraitis, V., Heath, A. C., Kubo, M., Larsson, A., Lindgren, C. M., Madden, P. A. F., Montgomery, G. W., Papanicolaou, G. J., Reiner, A. P., Sundstrom, J., Thornton, T. A., Lind, L., Ingelsson, E., Cai, J., Martin, N. G., Kooperberg, C., Matsuda, K., Whitfield, J. B., Okada, Y., Laurie, C. C., Morris, A. P., and Franceschini, N. (2016) Trans-ethnic fine mapping highlights kidney-function genes linked to salt sensitivity. Am J Hum Genet 99, 636–646

18. Robinson-Cohen, C., Lutsey, P. L., Kleber, M. E., Nielson, C. M., Mitchell, B. D., Bis, J. C., Eny, K. M., Portas, L., Eriksson, J., Lorentzon, M., Koller, D. L., Milaneschi, Y., Teumer, A., Pilz, S., Nethander, M., Selvin, E., Tang, W., Weng, L. C., Wong, H. S., Lai, D., Peacock, M., Hannemann, A., Volker, U., Homuth, G., Nauk, M., Murgia, F., Pattee, J. W., Orwoll, E., Zmuda, J. M., Riancho, J. A., Wolf, M., Williams, F., Penninx, B., Econs, M. J., Ryan, K. A., Ohlsson, C., Paterson, A. D., Psaty, B. M., Siscovick, D. S., Rotter, J. I., Pirastu, M., Streeten, E., Marz, W., Fox, C., Coresh, J., Wallaschofski, H., Pankow, J. S., De Boer, I. H., and Kestenbaum, B. (2017) Genetic variants associated with circulating parathyroid hormone. J Am Soc Nephrol 28, 1553–1565

19. Long, J., Chen, Y., Lin, H., Liao, M., Li, T., Tong, L., Wei, S., Xian, X., Zhu, J., Chen, J., Tian, J., Wang, Q., and Mo, Z. (2018) Significant association between RGS14 rs12654812 and nephrolithiasis risk among Guangxi population in China. J. Clin. Lab. Anal., e22435

20. Lek, M., Karczewski, K. J., Minikel, E. V., Samocha, K. E., Banks, E., Fennell, T., O’Donnell-Luria, A. H., Ware, J. S., Hill, A. J., Cummings, B. B., Tukiainen, T., Birnbaum, D. P., Kosmicki, J. A., Duncan, L. E., Estrada, K., Zhao, F., Zou, J., Pierce-Hoffman, E., Berghout, J., Cooper, D. N., Deflaux, N., DePristo, M., Do, R., Flannick, J., Fromer, M., Gauthier, L., Goldstein, J., Gupta, N., Howrigan, D., Kiezun, A., Kurki, M. I., Moonshine, A. L., Natarajan, P., Orozco, L., Peloso, G. M., Poplin, R., Rivas, M. A., Ruano-Rubio, V., Rose, S. A., Ruderfer, D. M., Shakir, K., Stenson, P. D., Stevens, C., Thomas, B. P., Tiao, G., Tusie-Luna, M. T., Weisburd, B., Won, H. H., Yu, D., Altshuler, D. M., Ardissino, D., Boehnke, M., Danesh, J., Donnelly, S., Elosua, R., Florez, J. C., Gabriel, S. B., Getz, G., Glatt, S. J., Hultman, C. M., Kathiresan, S., Laakso, M., McCarroll, S., McCarthy, M. I., McGovern, D., McPherson, R., Neale, B. M., Palotie, A., Purcell, S. M., Saleheen, D., Scharf, J. M., Sklar, P., Sullivan, P. F., Tuomilehto, J., Tsuang, M. T., Watkins, H. C., Wilson, J. G., Daly, M. J., MacArthur, D. G., and Exome Aggregation, C. (2016) Analysis of protein-coding genetic variation in 60,706 humans. Nature 536, 285–291

21. Squires, K. E., Montanez-Miranda, C., Pandya, R. R., Torres, M. P., and Hepler, J. R. (2018) Genetic analysis of rare human variants of regulators of g protein signaling proteins and their role in human physiology and disease. Pharmacol Rev 70, 446–474

22. Hung, A. Y., and Sheng, M. (2002) PDZ domains: structural modules for protein complex assembly. Journal of Biological Chemistry 277, 5699–5702

23. Romero, G., Von Zastrow, M., and Friedman, P. A. (2011) Role of PDZ proteins in regulating trafficking, signaling, and function of GPCRs. Means, motif, and opportunity. Advances in Pharmacology 62, 279–314

24. Beck, L., Karaplis, A. C., Amizuka, N., Hewson, A. S., Ozawa, H., and Tenenhouse, H. S. (1998) Targeted inactivation of *Npt2* in mice leads to severe renal phosphate wasting, hypercalciuria, and skeletal abnormalities. Proceedings of the National Academy of Sciences of the United States of America 95, 5372–5377

25. Weinman, E. J., Steplock, D., Shenolikar, S., and Biswas, R. (2011) Fibroblast growth factor-23-mediated inhibition of renal phosphate transport in mice requires sodium-hydrogen exchanger regulatory factor-1 (NHERF-1) and synergizes with parathyroid hormone. Journal of Biological Chemistry 286, 37216–37221

26. Shenolikar, S., Voltz, J. W., Minkoff, C. M., Wade, J. B., and Weinman, E. J. (2002) Targeted disruption of the mouse NHERF-1 gene promotes internalization of proximal tubule sodiumphosphate cotransporter type IIa and renal phosphate wasting. Proceedings of the National Academy of Sciences of the United States of America 99, 11470–11475

27. Karim, Z., Gerard, B., Bakouh, N., Alili, R., Leroy, C., Beck, L., Silve, C., Planelles, G., Urena-Torres, P., Grandchamp, B., Friedlander, G., and Prie, D. (2008) NHERF1 mutations and responsiveness of renal parathyroid hormone. New England Journal of Medicine 359, 1128–1135

28. Brone, B., and Eggermont, J. (2005) PDZ proteins retain and regulate membrane transporters in polarized epithelial cell membranes. American Journal of Physiology. Cell Physiology 288, C20–C29

29. Mamonova, T., Zhang, Q., Khajeh, J. A., Bu, Z., Bisello, A., and Friedman, P. A. (2015) Canonical and Noncanonical Sites Determine NPT2A Binding Selectivity to NHERF1 PDZ1. PLoS One 10, e0129554

30. Courbebaisse, M., Leroy, C., Bakouh, N., Salaun, C., Beck, L., Grandchamp, B., Planelles, G., Hall, R. A., Friedlander, G., and Prie, D. (2012) A new human NHERF1 mutation decreases renal phosphate transporter NPT2a expression by a PTH-independent mechanism. PLoS One 7, e34764

31. Biber, J., Malmström, K., Reshkin, S., and Murer, H. (1990) Phosphate transport in established renal epithelial cell lines. Methods in Enzymology 191, 494–504

32. Sneddon, W. B., Ruiz, G. W., Gallo, L. I., Xiao, K., Zhang, Q., Rbaibi, Y., Weisz, O. A., Apodaca, G. L., and Friedman, P. A. (2016) Convergent signaling pathways regulate parathyroid hormone and fibroblast growth factor-23 action on NPT2A-mediated phosphate transport. Journal of Biological Chemistry 291, 18632–18642

33. Emlet, D. R., Pastor-Soler, N., Marciszyn, A., Wen, X., Gomez, H., Humphries, W. H. t., Morrisroe, S., Volpe, J. K., and Kellum, J. A. (2017) Insulin-like growth factor binding protein 7 and tissue inhibitor of metalloproteinases-2: differential expression and secretion in human kidney tubule cells. Am J Physiol Renal Physiol 312, F284–F296

34. Wieser, M., Stadler, G., Jennings, P., Streubel, B., Pfaller, W., Ambros, P., Riedl, C., Katinger, H., Grillari, J., and Grillari-Voglauer, R. (2008) hTERT alone immortalizes epithelial cells of renal proximal tubules without changing their functional characteristics. Am J Physiol Renal Physiol 295, F1365–1375

35. Karthikeyan, S., Leung, T., and Ladias, J. A. (2002) Structural determinants of the Na^+^/H^+^ exchanger regulatory factor interaction with the ß_2_ adrenergic and platelet-derived growth factor receptors. J. Biol. Chem. 277, 18973–18978

36. Mamonova, T., Kurnikova, M., and Friedman, P. A. (2012) Structural basis for NHERF1 PDZ domain binding. Biochemistry 51, 3110–3120

37. Mamonova, T., Zhang, Q., Chandra, M., Collins, B. M., Sarfo, E., Bu, Z., Xiao, K., Bisello, A., and Friedman, P. A. (2017) Origins of PDZ Binding Specificity. A Computational and Experimental Study Using NHERF1 and the Parathyroid Hormone Receptor. Biochemistry 56, 2584–2593

38. Lamiable, A., Thevenet, P., Rey, J., Vavrusa, M., Derreumaux, P., and Tuffery, P. (2016) PEPFOLD3: faster de novo structure prediction for linear peptides in solution and in complex. Nucleic Acids Res 44, W449–454

39. Mahon, M. J., and Segre, G. V. (2004) Stimulation by parathyroid hormone of a NHERF-1-assembled complex consisting of the parathyroid hormone I receptor, PLCß, and actin increases intracellular calcium in opossum kidney cells. Journal of Biological Chemistry 279, 23550–23558

40. Wang, B., Yang, Y. M., and Friedman, P. A. (2008) Na/H exchange regulatory factor 1, a novel AKT-associating protein, regulates extracellular signal-regulated kinase signaling through a B-Raf-mediated pathway. Mol. Biol. Cell 19, 1637–1645

41. Khundmiri, S. J., Ahmad, A., Bennett, R. E., Weinman, E. J., Steplock, D., Cole, J., Baumann, P. D., Lewis, J., Singh, S., Clark, B. J., and Lederer, E. D. (2008) Novel regulatory function for NHERF-1 in Npt2a transcription. American Journal of Physiology. Renal Physiology 294, F840–F849

42. Cushing, P. R., Fellows, A., Villone, D., Boisguerin, P., and Madden, D. R. (2008) The relative binding affinities of PDZ partners for CFTR: a biochemical basis for efficient endocytic recycling. Biochemistry 47, 10084–10098

43. Mamonova, T., Zhang, Q., Chandra, M., Collins, B. M., Sarfo, E., Bu, Z., Xiao, K., Bisello, A., and Friedman, P. A. (2017) Origins of PDZ binding specificity. A computational and experimental case study using NHERF1 and the parathyroid hormone receptor. Biochemistry 56, 2584–2593

44. Clairfeuille, T., Mas, C., Chan, A. S., Yang, Z., Tello-Lafoz, M., Chandra, M., Widagdo, J., Kerr, M. C., Paul, B., Merida, I., Teasdale, R. D., Pavlos, N. J., Anggono, V., and Collins, B. M. (2016) A molecular code for endosomal recycling of phosphorylated cargos by the SNX27-retromer complex. Nat. Struct. Mol. Biol. 23, 921–932

45. Custer, M., Meier, F., Schlatter, E., Greger, R., Garcia-Perez, A., Biber, J., and Murer, H. (1993) Localization of NaPi-1, a Na-Pi cotransporter, in rabbit kidney proximal tubules--I. mRNA localization by reverse transcription/polymerase chain reaction. Pflügers Archiv. European Journal of Physiology 424, 203–209

46. Wang, B., Means, C. K., Yang, Y., Mamonova, T., Bisello, A., Altschuler, D. L., Scott, J. D., and Friedman, P. A. (2012) Ezrin-anchored PKA coordinates phosphorylation-dependent disassembly of a NHERF1 ternary complex to regulate hormone-sensitive phosphate transport. Journal of Biological Chemistry 287, 24148–24163

47. Cole, J. A., Forte, L. R., Krause, W. J., and Thorne, P. K. (1989) Clonal sublines that are morphologically and functionally distinct from parental OK cells. American Journal of Physiology 256, F672–F679

48. Case, D. A., Betz, R. M., Botello-Smith, W., Cerutti, D. S., Cheatham, T. E. I., Darden, T. A., Duke, R. E., Giese, T. J., Gohlke, H., Goetz, A. W., Homeyer, N., Izadi, S., Janowski, P., Kaus, J., Kovalenko, A., Lee, T. S., LeGrand, S., Li, P., Lin, C., Luchko, T., Luo, R., Madej, B., Mermelstein, D., Merz, K. M., Monard, G., Nguyen, H., Nguyen, H. T., Omelyan, I., Onufriev, A., Roe, D. R., Roitberg, A., Sagui, C., Simmerling, C. L., Swails, J., Walker, R. C., Wang, J., Wolf, R. M., Wu, X., Xiao, L., York, D. M., and Kollman, P. A. (2016) AMBER 16. University of California, San Francisco

